# Custom Probe-Based Spatial Transcriptomics Enables Microbiome Detection in FFPE Colorectal Cancer Tissue

**DOI:** 10.64898/2026.07.29.741338

**Authors:** Rayyan Aburajab, Jennifer L. Karmouch, Robert R. Jenq, David W. Craig

## Abstract

Intratumoral bacteria have emerged as functionally relevant components of the tumor microenvironment, yet the spatial relationship between these bacterial communities and host gene expression remains poorly characterized, in part due to methodological constraints. Existing spatial transcriptomics approaches for microbial detection rely on fresh frozen tissue, excluding FFPE specimens which dominate clinical archives. Here, we describe a custom probe design pipeline targeting the variable regions of bacterial 16S rRNA, compatible with the probe-based chemistry of the 10x Genomics Visium CytAssist platform, enabling spatially resolved bacterial profiling in FFPE tissue. Applied to a pilot cohort of six FFPE colorectal cancer tumor and normal adjacent tissue specimens, we show that integration of custom microbial probes into the Visium workflow preserves host transcriptomic structure, with clustering analysis recapitulating expected colonic cell type architecture. Bacterial signal was detected across all samples in a spatially patterned and focal manner, with one tumor sample exhibiting markedly elevated signal intensity and a distinct invasive distribution pattern, driven by spatially structured *Bacteroides-Phocaeicola* and *Porphyromonas* signals with divergent intratumoral trajectories. These findings establish the feasibility of probe-based spatial metatranscriptomics in FFPE tissue and provide a generalizable framework for studying host-microbiome interactions in relevant clinical samples.

## INTRODUCTION

The tumor microbiome is increasingly recognized as an important component of the tumor microenvironment, with intratumoral bacteria shown to influence tumor progression,^1,2^ immune modulation,^3^ and even treatment response.^4^ These effects occur through diverse mechanisms, including direct DNA damage and genotoxicity,^5^ activation of oncogenic signaling pathways,^6^ promotion of chronic inflammation,^7^ and immune evasion.^8^ Much of what is known has emerged from studies centered on individual tumor-enriched species with distinct pro-tumorigenic mechanisms, such as *Fusobacterium nucleatum* in colorectal cancer (CRC),^7^ *Helicobacter pylori* in gastric cancer,^6^ *Bacteroides fragilis* in breast cancer,^9^ and *Porphyromonas gingivalis* in lung cancer.^10^ However, this species-specific lens likely oversimplifies the broader picture, as the collective composition and spatial localization of bacterial communities within the tumor microenvironment are increasingly thought to shape their functional impact.^11,12^

Characterizing how bacterial communities interact with host cells in their spatial context requires methods that can resolve both microbial identity and host gene expression within intact tissue. Existing approaches each address part of this problem but fall short of the full picture. While microbial sequencing technologies, such as 16S rRNA sequencing^13^ and whole metagenome shotgun sequencing,^14^ can provide compositional profiles of microbial communities, they are limited to bulk tissue analysis and lack the cellular and spatial resolution necessary to establish definitive associations with host cell states or tissue compartments. In contrast, in situ hybridization technologies like FISH^15^ and RNAscope^16^ can localize specific microbial targets within tissue sections,^11^ but are limited by low throughput and multiplexing capacity, which precludes broad community- and host-wide profiling.

Spatial transcriptomics (ST) can address these limitations by enabling simultaneous detection of microbial signal and host gene expression across intact tissue sections. Recent studies have adapted the 10x Genomics Visium platform for bacterial detection by exploiting non-specific hybridization of 16S rRNA to poly(dT) capture probes,^11^ by incorporating universal 16S rRNA probes alongside the poly(dT) probes,^17^ or by enzymatically polyadenylating bacterial RNA in situ to enable their capture by poly(dT) probes.^18^ These approaches have collectively established that spatial co-profiling of microbial communities and host transcriptomes is achievable, revealing spatially structured host-microbe interactions that bulk methods cannot resolve. However, while innovative, all of these strategies are constrained by their reliance on the Visium v1 platform, which is compatible only with fresh frozen tissue, which typically requires prospective collection and represents a small fraction of available clinical material. Formalin-fixed paraffin-embedded (FFPE) tissue, by contrast, dominates clinical archives and is far more abundant for retrospective studies, yet no validated method currently exists for spatially resolved microbial profile in this format. This represents a fundamental limitation in the field, restricting spatial microbiome methods to prospectively collected samples and excluding the vast archives of available FFPE tissue, thus limiting broader clinical applications.

The 10x Genomics Visium CytAssist Gene Expression platform (Visium v2), on the other hand, was designed to extend ST to FFPE tissue, broadening its applicability to clinically archived samples. Unlike Visium v1, this approach uses a targeted, probe-based chemistry in which target-specific pre-designed oligonucleotide probe pairs hybridize to their target transcripts and are themselves sequenced, such that target identity is determined entirely by the probe sequence. Adapting this platform for bacterial detection would require designing and integrating taxon-specific probes alongside the standard human transcriptome panel. To our knowledge, such an approach has not been previously demonstrated. Targeting the variable regions of bacterial 16S rRNA, the standard marker for bacterial taxonomic identification,^19^ would be well-suited for this purpose. CRC represents one of the most extensively studied contexts for tumor-associated bacteria, harboring a well-characterized and microbe-rich microenvironment,^20^ making it an ideal setting in which to develop and apply new methods for spatial microbial profiling.

Here, we describe generating a custom 16S probe design pipeline and resulting probe pool targeting a broad panel of bacterial taxa relevant to the CRC-associated microbiome. In a pilot cohort composed of CRC as well as histologically normal tissues adjacent tissue (NAT) samples, we demonstrate that microbial signal is detectable, spatially patterned, and biologically coherent. Our study establishes the feasibility of probe-based spatial meta-transcriptomics in FFPE tissue and provides a framework that can be adapted to other microbial targets and tissue contexts.

## RESULTS

### Development of a Custom Microbial Probe Panel for Visium FFPE

To determine the feasibility of profiling the microbiome in FFPE tissue using a probe-based sequencing spatial transcriptomics approach, we first asked whether bacterial 16S rRNA could serve as a viable probe target within the constraints of the Visium CytAssist platform, which requires probes to be designed as paired oligonucleotides with specific architectural requirements (Figure 1a, Methods). Using a custom computational pipeline (Figure 1b, Methods), we successfully designed a panel of platform-compatible probe pairs targeting bacterial taxa with established or emerging roles in the CRC-associated microbiome, including both CRC-enriched and CRC-depleted species (Supplementary Table 1). The resulting probe set, containing 195 probe pairs targeting 15 genera and 4 families (Table 1), was synthesized and spiked into the standard Visium human transcriptome probe set (Figure 1c). This modified panel was applied to a pilot cohort of six FFPE colorectal specimens (Table 2, Methods), comprising three CRC tumor and three normal adjacent tissue (NAT) tissue, processed using the standard Visium FFPE workflow (Figure 1d-f). Across all samples, both host transcriptomic and bacterial signals were detected, establishing that microbial probes can be successfully integrated into the Visium CytAssist workflow.

**Table 1.**
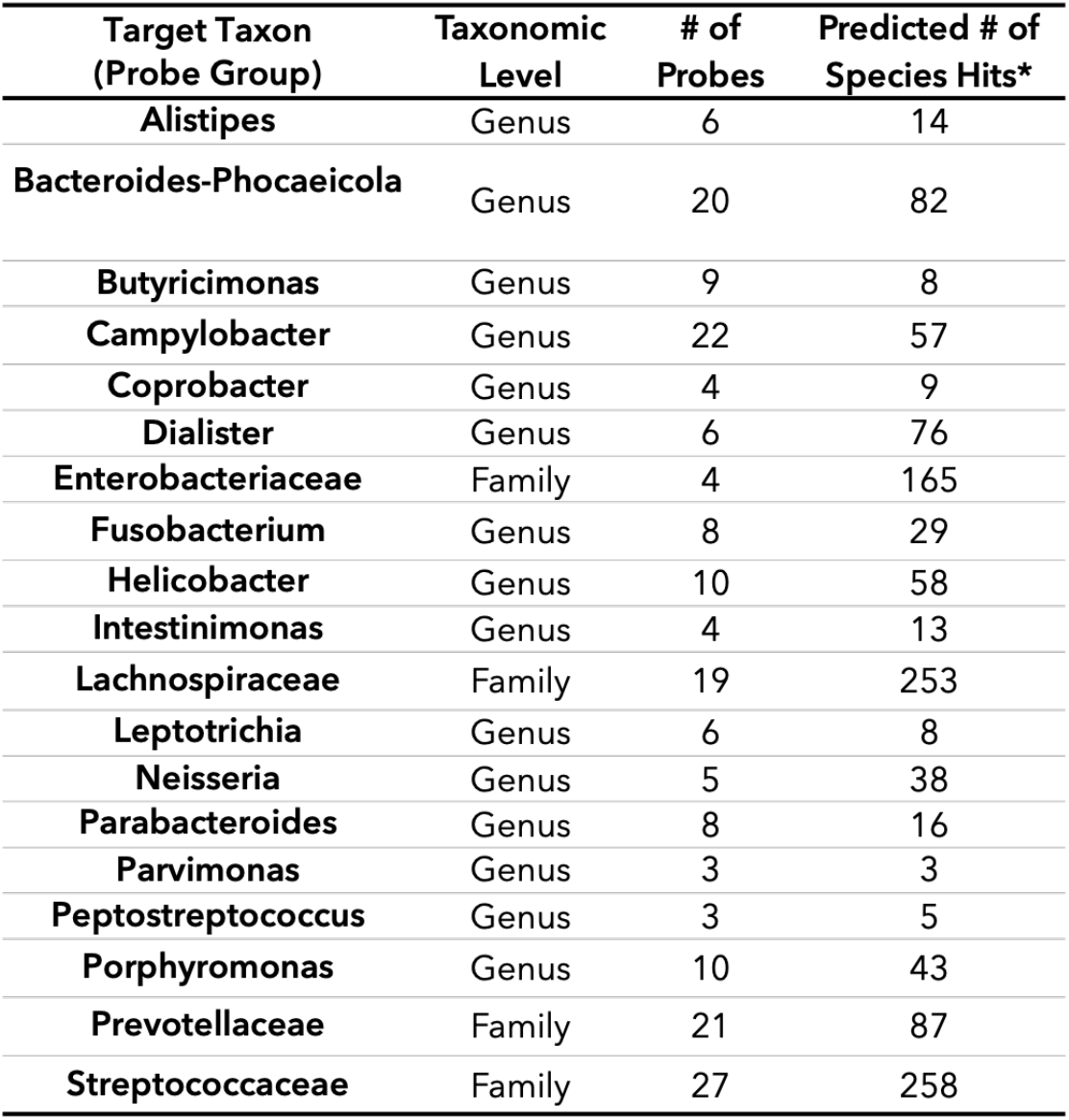
Composition of the custom probe panel. Probe taxon-level specificity, probe count per group, and the predicted number of species potentially detectable per probe group for each target taxon. * Based on NCBI BLAST percent identity ≥ 90 and query coverage ≥ 0.9

**Table 2.** Clinical and histopathological characteristics of the FFPE colorectal tissue specimens.

| Sample ID | Type | Age | Gender | Stage | Grade | Tumor Area (%) |
| --- | --- | --- | --- | --- | --- | --- |
| **NAT** |  |  |  |  |  |  |
| CRC 03 | Colon Normal/Benign | 52 | Female | - | - | - |
| CRC 06 | Colon Normal/Benign | 48 | Male | - | - | - |
| CRC 08 | Colon Normal/Benign | 63 | Female | - | - | - |
| **CRC** |  |  |  |  |  |  |
| CRC 03 | Colorectal Cancer, Adenocarcinoma | 52 | Female | III-B | G2 | 30% |
| CRC 07 | Colorectal Cancer, Adenocarcinoma | 45 | Female | III-B | G3 | 40% |
| CRC 08 | Colorectal Cancer, Adenocarcinoma | 63 | Female | II-A | low | 73% |

Includes sample type, patient age, gender, tumor stage and grade, and estimated tumor cellularity.

**Figure 1.**
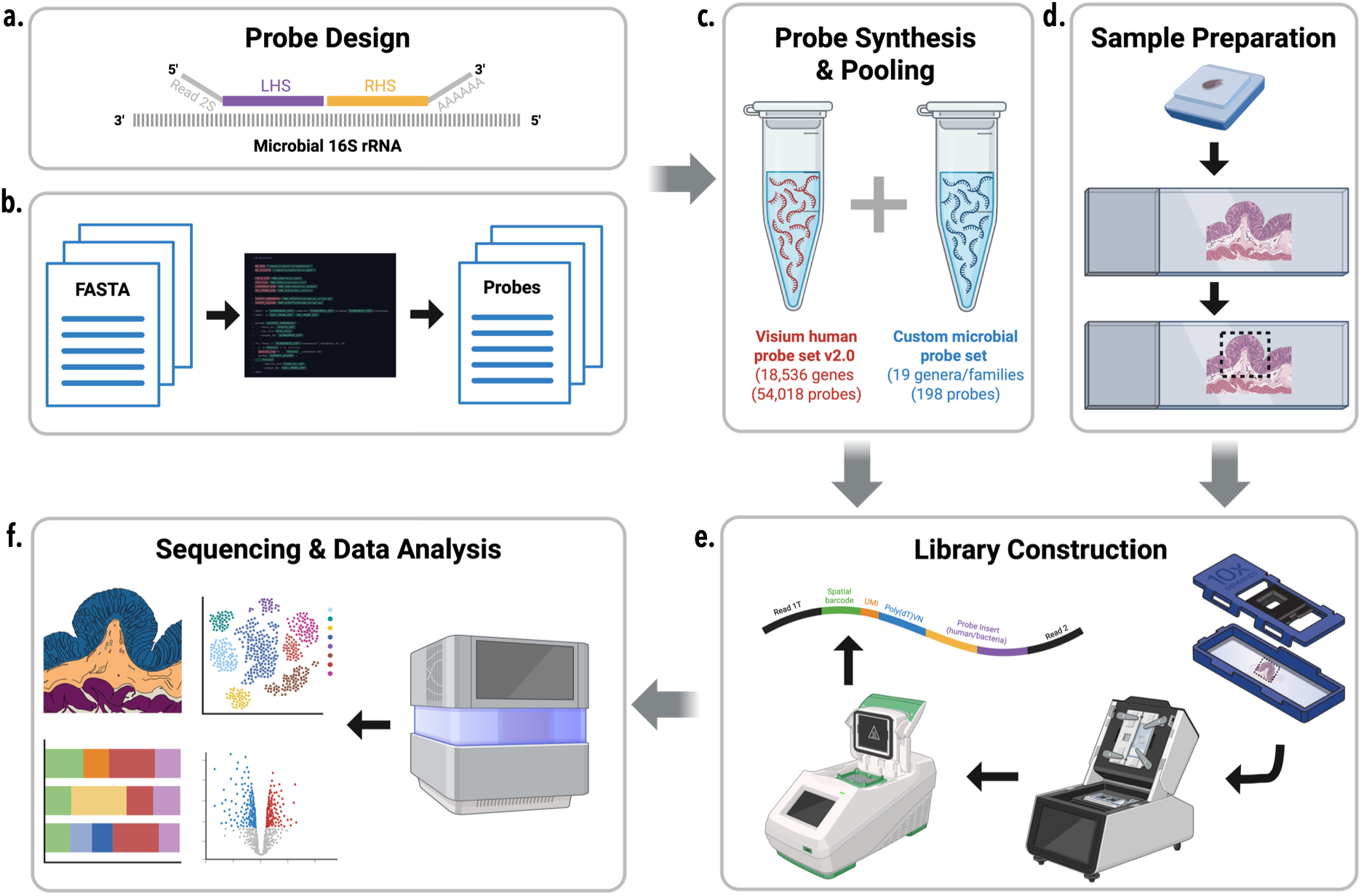
Overview of microbiome probe design and ST workflow. **(a)** Following 10x Genomics guidelines, probes are designed for split-probe ligation chemistry using LHS/RHS probes. **(b)** Target microbial 16S rRNA sequences are provided as NCBI RefSeq FASTA files and used to generate candidate probes, which are filtered and quality controlled for optimal taxonomic specificity and performance. **(c)** The resulting microbial probe set is synthesized and spiked into the Visium human transcriptome probe set to create a combined probe pool. **(d)** FFPE tissue blocks are sectioned, stained with H&E, and ROIs are selected for capture. **(e)** The combined probe pool is hybridized to tissue sections, followed by CytAssist-mediated transfer and library preparation. **(f)** Sequencing libraries are processed and analyzed to generate spatially resolved transcriptomic and microbial profiles.

### Initial Visium Results Demonstrate Preservation of Host Transcriptomic Structure

To confirm that inclusion of custom microbial probes did not disrupt host transcriptomic structure, we performed unsupervised clustering across the integrated sample set following batch correction and dimensionality reduction. UMAP visualization revealed well-defined clusters and shared transcriptional structure across all six colonic samples (Figure 2a). Examination of per-cluster sample composition revealed that some clusters broadly segregated into those enriched for normal, tumor or sample pair-specific contributions, reflecting both tissue type and sample-level transcriptional variation (Figure 2b). These results indicate that the bacterial probe-derived reads did not compromise the integrity of host gene expression profiling.

**Figure 2.**
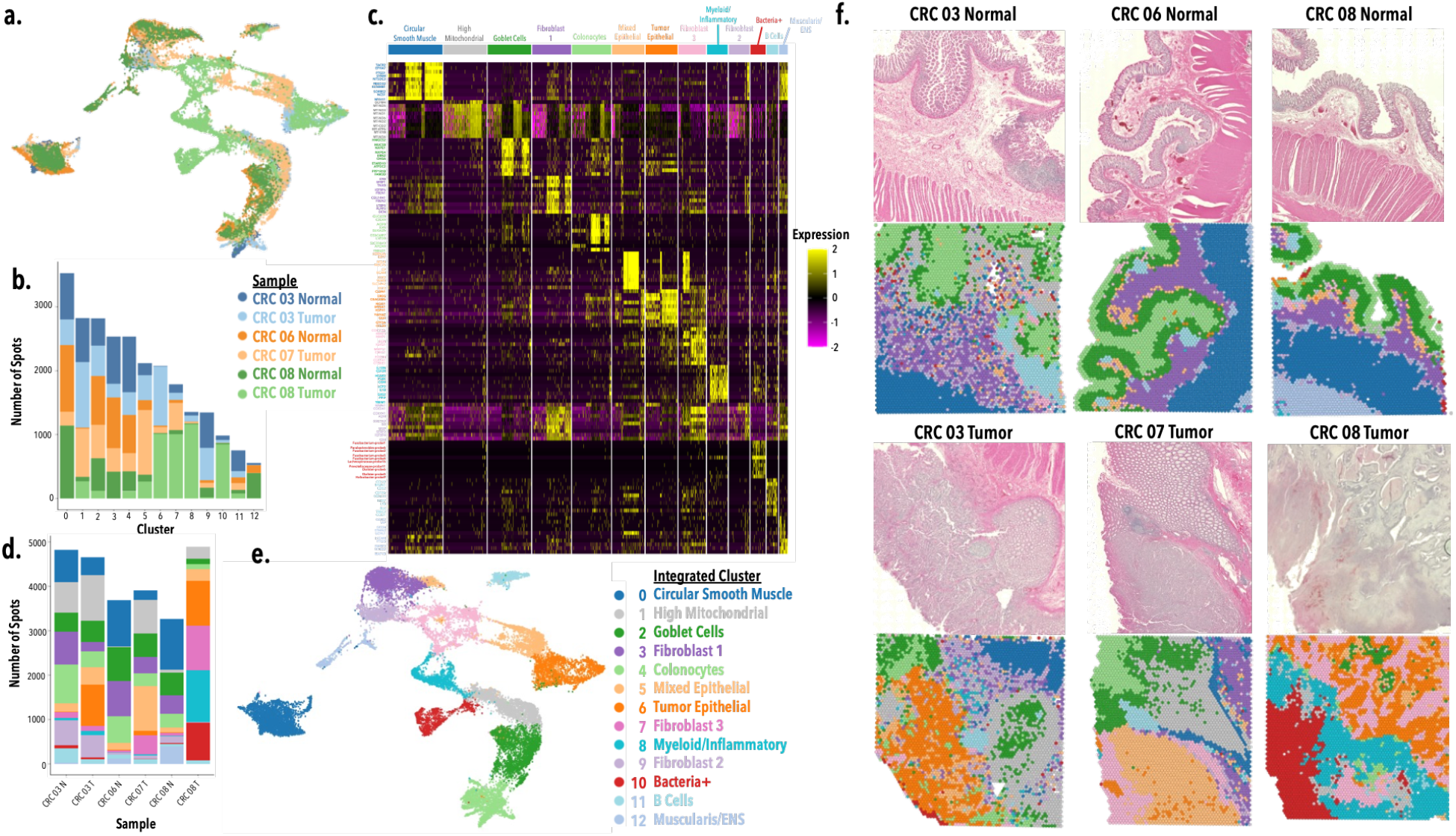
Integrative ST analysis of colorectal tissue reveals preservation of host transcriptomic structure. **(a)** UMAP visualization of all six samples following SCTransform normalization, Harmony batch correction, and unsupervised Louvain clustering, colored by sample identity **(b)** Per-cluster sample composition by tissue type. **(c)** Heatmap of cluster marker gene expression used for cell type annotation. **(d)** Cell-type composition across individual samples. **(e)** UMAP colored by annotated cell type identity. **(f)** H&E images and corresponding spatial projection of annotated cell types onto tissue sections (NAT, upper; CRC tumor, lower).

To characterize the transcriptional identity of each cluster, differential expression analysis was performed, revealing gene signatures consistent with major colonic cell populations. These signatures enabled cell type annotation across the integrated dataset (Figure 2c, e), the composition of which across individual samples is summarized in Figure 2d. Projection of annotated clusters into tissue space (Figure 2f) revealed spatial domains that aligned with expected layered architecture of colon tissue. Epithelial clusters included colonocytes (GUCA2B, AQP8, CEACAM7) and goblet cells (MUC5B, CHGA), both predominantly enriched in NAT samples and reflective of normal mucosal differentiation. Two epithelial clusters with tumor-associated signatures were also identified: a mixed epithelial cluster (CDKN2A, KRT7, MSLN) predominantly derived from tumor samples with minor normal contributions, and a tumor epithelial cluster (EREG, CEACAM6, SPINK1) composed almost exclusively of malignant tissue. The muscularis propria was represented by a cluster expressing smooth muscle markers (MYH11, SORBS2, TACR2) corresponding to the circular muscle layer, while a spatially adjacent cluster expressing enteric nervous system markers (UCHL1, L1CAM, STMN2, SCGN) likely corresponds to the myenteric plexus, though spatial overlap with the longitudinal muscle layer was observed in some samples. Three fibroblast populations were also identified. Fibroblast 1 (DCN, FBLN1, FBLN2, TNXB) and Fibroblast 2 (COL1A1, VIM) showed expression profiles and spatial localization consistent with submucosal fibroblasts, with Fibroblast 1 predominantly enriched in NAT samples and Fibroblast 2 largely contributed by CRC 03 specimens. Fibroblast 3 (MMP1, MMP3, COL11A1, POSTN) expressed markers associated with tumor stromal remodeling, was predominantly derived from tumor samples, and was spatially distinct from the other fibroblast populations. Immune populations included a myeloid/inflammatory cluster (IL1B, TREM1, CSF3R), which formed a prominent region in CRC 08 Tumor with sparser representation across the remaining samples, and a B cell cluster (MS4A1, CD79A, CD52) presenting as lymphoid follicle-like aggregates across both normal and tumor specimens. A distinct cluster characterized by high bacterial probe signal was identified, with top markers corresponding to probes targeting CRC-relevant taxa, including *Fusobacterium* and *Lachnospiraceae*, and was transcriptionally isolated from host cell populations. A cluster marked by high mitochondrial gene expression was observed across samples at varying tissue locations, likely corresponding to regions of low molecular diversity. Together, these clusters reflect the expected cellular architecture of colonic tissues across both normal and tumor compartments.

### Integration of Custom Probes Enables Spatial Microbial Signal Detection

Building on these findings, we next examined the spatial distribution of microbial signal across tissues. First, spatial projection of total bacteria probe reads (log1p-transformed counts) revealed detectable microbial signal across samples (Figure 3a). As indicated by white arrows, signal was focally enriched in spatial hotspots rather than uniformly distributed, with marked variation in signal intensity between samples. In NAT samples (Figure 3a, upper), microbial signal was primarily confined to the luminal surface when present (white arrows), consistent with the known organization of the normal colonic microbiome, in which bacteria are confined to the mucus surface layer. The overall signal intensity is consistent with expected clinical specimen collection and processing conditions. In tumor samples (Figure 3a, lower), signal was similarly focal in CRC 03 and CRC 07 (white arrows), while CRC 08 showed substantially greater signal intensity and deeper spatial distribution throughout the tissue, representing the most bacterially enriched sample in the cohort. Next, examination of the relative contribution of each bacterial group across samples revealed marked compositional differences (Figure 3b). In samples with low overall bacterial signal, compositional profiles were dominated by *Streptococcaceae*; however, given the low absolute signal and non-localized pattern in these samples, this likely reflects background rather than true microbial enrichment.

**Figure 3.**
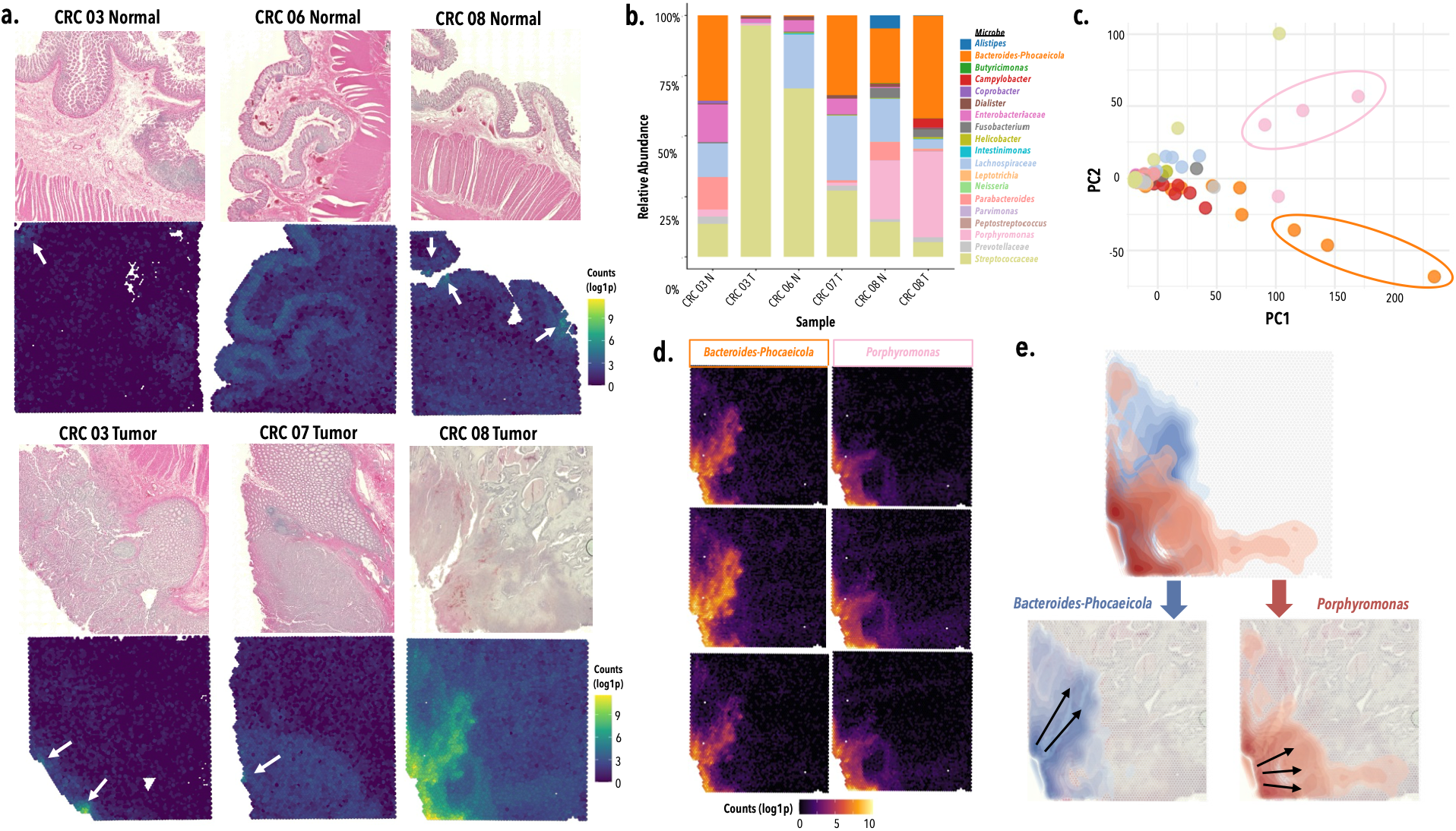
Custom microbial probes enable spatially resolved detection of bacteria in colorectal tissue. H&E images and corresponding spatial projection of total bacterial probe counts (log1p-transformed) onto samples (NAT, upper; CRC tumor, lower). White arrows indicate focal regions of microbial signal enrichment. Stacked bar plot showing the relative compositional contribution of each bacterial group across samples. PCA of the CLR-transformed probe-level count matrix, with probes plotted in PC1-PC2 space. Three *Bacteroides-Phocaeicola* and three *Porphyromonas* are circled. **(d)** Spatial projection of probe counts (log1p-transformed) for the six PCA-identified probes onto CRC 08 Tumor tissue, grouped by *Bacteroides-Phocaeicola* (left) and *Porphyromonas* (right). **(e)** Heatmap contour plots showing the spatial density of PCA-identified *Bacteroides-Phocaeicola* and *Porphyromonas* signal within CRC 08 Tumor tissue (combined, upper; separated, lower). Arrows indicate the directionality of inward signal gradients.

To assess whether distinct bacterial subpopulations could be distinguished across the cohort, we performed PCA on the CLR-transformed probe-level count matrix, following filtering of low detection spots and sparsely detected probes. Visualization of the first two principal components identified two distinct probe clusters, with three *Porphyromonas* probes driving PC1 and three *Bacteroides-Phocaeicola* probes driving PC2 (Figure 3c). Spatial visualization of these six probes in the CRC 08 Tumor sample revealed that the *Porphyromonas* probe group and the *Bacteroides-Phocaeicola* probe group each displayed highly consistent spatial patterns with one another (Figure 3d), suggesting coherent probe group-level signal. Since the spatial distributions of the two groups appeared visually distinct, we next characterized their density patterns using a heatmap contour plot (Figure 3e). Examination of the spatial density of each group revealed a shared gradient structure, with signal concentrated at the tissue edge and decreasing inward, suggestive of an invasive distribution pattern. Notably, *Porphyromonas* and *Bacteroides-Phocaeicola* displayed distinct inward gradient trajectories, suggesting these taxa may be infiltrating the tumor tissue along different spatial axes. Together, these findings demonstrate the ability of the custom microbial probe pool to detect spatially resolved bacterial signal with sufficient signal resolution to distinguish distinct tissue distribution patterns of co-occurring bacterial taxa within CRC tumor tissue.

## DISCUSSION

In this study, we developed and evaluated a custom microbial probe pool compatible with the Visium CytAssist platform, demonstrating the feasibility of spatially resolved microbiome profiling in FFPE colorectal tissue. Integration of the microbial probe pool into the standard Visium workflow did not compromise host transcriptomic data quality, with downstream analysis yielding well-defined transcriptional clusters and spatially coherent cell type distributions consistent with expected colonic tissue architecture. Bacterial signal was detected across all samples, though predominantly at low levels with focal enrichment in discrete hotspots. The low signal observed across NAT and low-signal tumor samples is consistent with expectations for human-derived clinical specimens.

CRC 08 emerged as a compositionally distinct sample, with PCA of the probe-level count matrix identifying *Bacteroides-Phocaeicola* and *Porphyromonas* as the primary drivers of microbial signal, both displaying spatially structured and directionally distinct distribution patterns within the tumor. The mechanistic diversity through which these two taxa interact with the tumor microenvironment may underlie the divergent spatial distribution patterns observed here. The *Bacteroides* genus is among the most abundant commensal taxa in the human gut microbiome, with certain species recognized as opportunistic pathogens across various gastrointestinal diseases.^21^ In the context of CRC, enterotoxigenic *Bacteroides fragilis* (ETBF) represents a well-characterized example, which secretes a metalloprotease toxin that disrupts epithelial barrier integrity and activates STAT3-mediated inflammatory signaling to promote tumorigenesis.^22^ Similarly, members of the *Porphyromonas* genus include species with established roles in CRC tumorigenesis. For example, *Porphyromonas gingivalis* has been implicated through both direct tumor-promoting and immunomodulatory mechanisms, including tissue invasion and butyrate-mediated induction of cellular senescence,^23^ activation of proliferative signaling pathways,^24^ NLRP3 inflammasome-driven inflammation,^25^ and immune evasion.^26^ Whether the distinct distribution patterns reflect biologically meaningful differences in colonization behavior of these taxa will require further investigation in larger cohorts in addition to orthogonal validation through other methods, like bulk 16S rRNA sequencing or RNAscope-based in situ hybridization, to further strengthen confidence in the detected signal.

While these findings demonstrate the feasibility of spatially resolved microbiome profiling in FFPE tissue, several methodological constraints should be considered. As a targeted approach, a fundamental limitation of this method is that targets must be selected in advance, precluding discovery-based characterization of in-situ microbial communities. However, the Visium platform supports substantially larger custom probe panels within the existing workflow guidelines,^27^ leaving considerable capacity for both targeted expansion and the inclusion of controls beyond the scope used here. For instance, the incorporation of universal bacterial probes could complement the targeted panel by providing an estimate of total bacterial load independent of the selected taxa, and probes targeting common laboratory bacterial contaminants could be included to account for potential background signal. The spatial nature of this platform does, however, offers some inherent protection against contamination, as contaminant signal would be expected to appear diffusely rather than in structured spatial patterns. Probe specificity is a particular consideration for microbial 16S rRNA targets, which contain regions of both high conservation and sequence variability, and differ substantially from the RNA targets for which the 10x Genomics custom probe guidelines were developed. NCBI BLAST filtering thresholds of percent identity ≥90 and query coverage ≥0.9 were applied to approximate these specificity criteria, though further optimization may be warranted. Variations in tissue processing conditions inherent to archival-based studies may introduce variability in nucleic acid preservation and signal detection. Standard FFPE processing is known to compromise mucus layer integrity,^28^ and pre-surgical bowel preparation, including mechanical preparation and routinely administered antibiotic prophylaxis, substantially reduces bacterial load in resected specimens.^29^ These factors are expected to reduce surface-associated bacterial signal and should be considered when interpreting microbiome profiles derived from surgically resected colorectal FFPE tissue. However, those concerns are of less concern given that this approach is designed to profile the intratumoral microbiome rather than luminal or mucosal communities. Careful consideration of these constraints, along with further optimization of probe design and panel composition, will be important for future applications of this approach. Together, these findings establish a framework for spatially resolved microbial profiling in FFPE tissue that can be adapted to other microbial targets and tissue contexts.

## METHODS

### Sample Selection

Human-derived FFPE colorectal tissue blocks were purchased from Discovery Life Sciences. RNA was extracted from tissue scrolls using the Qiagen AllPrep DNA/RNA FFPE kit following the standard manufacturer protocol (Qiagen #80234) and RNA quality was assessed by DV200 analysis on the Agilent TapeStation. Six samples with DV200 ≥ 30% were selected for downstream analysis. The cohort comprised three primary CRC tumor and three NAT specimens, with two patients contributing matched CRC-NAT pairs and the remaining two samples unmatched. Clinical and histopathological characteristics of all samples are summarized in Table 2.

### Probe Design

A custom probe pool was designed targeting distinct regions of microbial 16S rRNA, constructed as paired oligonucleotides analogous to the LHS/RHS probe architecture of the Visium CytAssist Gene Expression platform. Target microbial taxa were selected based on their relevance to the human oral and gut microbiomes, encompassing species with established or emerging roles in the CRC-associated microbiome based on publicly available pan-cancer and CRC-specific microbiome datasets, including The Cancer Microbiome Atlas (TCMA)^30^ and the Human Gut Microbiome Atlas (HGMA).^31^ To ensure broad detection of probes across taxonomic targets and to accommodate the technical constraints of Visium probe design, probes were designed at the genus or family level rather than the species level. Full length 16S rRNA sequences for each target species were obtained from the NCBI RefSeq 16S rRNA database (Supplementary Table 1), with consensus sequences generated using MAFFT multiple sequence alignment for species represented by multiple strains. Custom Python scripts were used to generate 50 bp probe pairs (LHS, RHS; 25 bp each), filtered according to 10x Genomics probe design criteria (Technical Note CG000621) including GC content (44−72%), absence of homopolymers (<4 repeats), and a thymine residue at position 25. Probe specificity was assessed by BLAST search against the NCBI nucleotide database, retaining only probes with percent identity ≥ 90 and query coverage ≥ 0.9 to the intended target taxon. Retained probes underwent secondary quality control analysis to evaluate melting temperature and predicted secondary structure. The resulting probe set comprised 195 probe pairs targeting 15 genera and 4 families, with 3 to 27 probes per taxon depending on sequence diversity. Probes targeting the human housekeeping gene GAPDH were included as positive controls for probe design integrity. The final probe pool was synthesized as an oligonucleotide pool at 50 pmol per oligo scale (Integrated DNA Technologies).

### Sequencing Library Preparation

FFPE colorectal tissue blocks were section at 10 μm thickness and subsequently processed using the Visium CytAssist Gene Expression platform (10x Genomics #1000443) following the standard manufacturer protocol for FFPE tissue. The custom microbial probe pool was incorporated into the hybridization mix alongside the standard Visium Human Whole Transcriptome Probe Set (v2.0), with each custom probe at a final concentration of 24 nM (LHS and RHS combined), per 10x Genomics guidelines for custom probe integration (Technical Note CG000621). Sequencing was performed on the NextSeq 2000 platform (Illumina) using XLEAP-SBS P2 reagents with a 100-cycle configuration.

### Data pre-processing

Raw sequencing data were processed using SpaceRanger (v3.1.3). To enable simultaneous quantification of host transcripts and microbial signal, a custom reference was constructed using the `spaceranger mkref` command. Microbial probe target sequences were appended as artificial chromosomes to the GRCh38 reference genome (refdata-gex-GRCh38-2020-A) fasta and GTF annotation files, generating a modified reference (refdata-gex-GRCh38-Microbiome-v1). Each sample was processed using the `spaceranger count` command with the modified reference supplied using the `–transcriptome` flag and the human transcriptome probe set (v2.0) file appended with the custom microbial probes supplied using the `--probe-set` flag.

### Data Analysis

All analyses were performed in R (v4.4.1) using Seurat (v5.2.0). Spots with low transcript counts were filtered based on tissue quality. Samples were normalized using `SCTransform`, merged into a single Seurat object, and PCA was performed on the top 5000 highly variable genes using 50 principal components and batch correction was performed using Harmony. The `FindNeighbors` function was applied using the first 30 dimensions, followed by unsupervised clustering (0.2 resolution) and UMAP visualization. Cluster marker genes were identified using `FindAllMarkers`, filtered by adjusted p-value (<0.05), pct.1 (>0.1), and log2 fold change (≥0.5), with the top 10 markers per cluster selected by average log2 fold change for heatmap visualization and cluster annotation.

### Microbial Signal Analysis

Bacterial probe counts were extracted from the normalized Seurat object. Spot-level quality control was applied to retain spots with non-zero total bacterial counts and a second highest normalized probe count greater than 2, ensuring detectable signal across at least two probes. Probe-level quality control was subsequently applied to retain probes detected above this threshold in a minimum of 4 spots, and spots with no remaining probe signal after probe filtering were removed. For PCA of the cohort-wide microbial probe matrix, CLR transformation was applied to raw probe counts with a pseudocount of 0.5 and PCA was performed using `prcomp` with unit variance scaling, with probes as observations. Probe scores across PC1 and PC2 were used to identify probes driving sample separation. For spatial density visualization, raw counts for the identified probes were summed per taxonomic group and log1p-transformed. Spots with values greater than 2 were retained to reduce background noise, and retained spots were weighted by their expression values prior to two-dimensional kernel density estimation using `stat_density_2d` in ggplot2.

The probe design pipeline and all analysis scripts are publicly available at: https://github.com/COHCCC/aburajab-spatial-microbiome-crc-2025

## Supporting information

Supplementary Probe Sequences

## Notes

https://www.ncbi.nlm.nih.gov/geo/query/acc.cgi?acc=GSE334323

https://github.com/COHCCC/aburajab-spatial-microbiome-crc-2025

### Competing Interest Statement

The authors have declared no competing interest.

## REFERENCES

1 Fu, A. et al. Tumor-resident intracellular microbiota promotes metastatic colonization in breast cancer. Cell 185, 1356–1372.e1326 (2022). 10.1016/j.cell.2022.02.027

2 Jin, C. et al. Commensal Microbiota Promote Lung Cancer Development via γδ T Cells. Cell 176, 998–1013.e1016 (2019). 10.1016/j.cell.2018.12.040

3 Kostic, A. D. et al. Fusobacterium nucleatum potentiates intestinal tumorigenesis and modulates the tumor-immune microenvironment. Cell Host Microbe 14, 207–215 (2013). 10.1016/j.chom.2013.07.007

4 Yu, T. et al. Fusobacterium nucleatum Promotes Chemoresistance to Colorectal Cancer by Modulating Autophagy. Cell 170, 548–563.e516 (2017). 10.1016/j.cell.2017.07.008

5 Wilson, M. R. et al. The human gut bacterial genotoxin colibactin alkylates DNA. Science 363 (2019). 10.1126/science.aar7785

6 Buti, L. et al. Helicobacter pylori cytotoxin-associated gene A (CagA) subverts the apoptosis-stimulating protein of p53 (ASPP2) tumor suppressor pathway of the host. Proc Natl Acad Sci U S A 108, 9238–9243 (2011). 10.1073/pnas.1106200108

7 Rubinstein, Mara R. et al. Fusobacterium nucleatum Promotes Colorectal Carcinogenesis by Modulating E-Cadherin/β-Catenin Signaling via its FadA Adhesin. Cell Host & Microbe 14, 195–206 (2013). 10.1016/j.chom.2013.07.012

8 Borowsky, J. et al. Association of Fusobacterium nucleatum with Specific T-cell Subsets in the Colorectal Carcinoma Microenvironment. Clinical Cancer Research 27, 2816–2826 (2021). 10.1158/1078-0432.Ccr-20-4009

9 Parida, S. et al. A Procarcinogenic Colon Microbe Promotes Breast Tumorigenesis and Metastatic Progression and Concomitantly Activates Notch and β-Catenin Axes. Cancer Discov 11, 1138–1157 (2021). 10.1158/2159-8290.Cd-20-0537

10 Liu, Y. et al. Clinical significance and prognostic value of Porphyromonas gingivalis infection in lung cancer. Transl Oncol 14, 100972 (2021). 10.1016/j.tranon.2020.100972

11 Galeano Niño, J. L. et al. Edect of the intratumoral microbiota on spatial and cellular heterogeneity in cancer. Nature 611, 810–817 (2022). 10.1038/s41586-022-05435-0

12 Nejman, D. et al. The human tumor microbiome is composed of tumor type–specific intracellular bacteria. Science 368, 973–980 (2020). 10.1126/science.aay9189

13 Caporaso, J. G. et al. Global patterns of 16S rRNA diversity at a depth of millions of sequences per sample. Proceedings of the National Academy of Sciences 108, 4516–4522 (2011). 10.1073/pnas.1000080107

14 Quince, C., Walker, A. W., Simpson, J. T., Loman, N. J. & Segata, N. Shotgun metagenomics, from sampling to analysis. Nature Biotechnology 35, 833–844 (2017). 10.1038/nbt.3935

15 Lukumbuzya, M., Schmid, M., Pjevac, P. & Daims, H. A Multicolor Fluorescence in situ Hybridization Approach Using an Extended Set of Fluorophores to Visualize Microorganisms. Frontiers in Microbiology Volume 10 - 2019 (2019). 10.3389/fmicb.2019.01383

16 Wang, F. et al. RNAscope: a novel in situ RNA analysis platform for formalin-fixed, paradin-embedded tissues. J Mol Diagn 14, 22–29 (2012). 10.1016/j.jmoldx.2011.08.002

17 Saarenpää, S. et al. Spatial metatranscriptomics resolves host–bacteria–fungi interactomes. Nature Biotechnology 42, 1384–1393 (2024). 10.1038/s41587-023-01979-2

18 Ntekas, I. et al. Spatial transcriptomics maps host–gut microbiome biogeography at high resolution. Nature Microbiology (2026). 10.1038/s41564-026-02286-7

19 Clarridge, J. E., 3rd. Impact of 16S rRNA gene sequence analysis for identification of bacteria on clinical microbiology and infectious diseases. Clin Microbiol Rev 17, 840–862, table of contents (2004). 10.1128/cmr.17.4.840-862.2004

20 Fan, Y. et al. Intratumoral microbiome: a crucial regulating factor in development and progression of colorectal cancer. Mol Biomed 6, 138 (2025). 10.1186/s43556-025-00376-2

21 Shin, J. H. et al. Bacteroides and related species: The keystone taxa of the human gut microbiota. Anaerobe 85, 102819 (2024). 10.1016/j.anaerobe.2024.102819

22 Chung, L. et al. Bacteroides fragilis Toxin Coordinates a Pro-carcinogenic In?ammatory Cascade via Targeting of Colonic Epithelial Cells. Cell Host & Microbe 23, 203–214.e205 (2018). 10.1016/j.chom.2018.01.007

23 Okumura, S. et al. Gut bacteria identified in colorectal cancer patients promote tumourigenesis via butyrate secretion. Nature Communications 12, 5674 (2021). 10.1038/s41467-021-25965-x

24 Mu, W. et al. Intracellular Porphyromonas gingivalis Promotes the Proliferation of Colorectal Cancer Cells via the MAPK/ERK Signaling Pathway. Front Cell Infect Microbiol 10, 584798 (2020). 10.3389/fcimb.2020.584798

25 Wang, X. et al. Porphyromonas gingivalis Promotes Colorectal Carcinoma by Activating the Hematopoietic NLRP3 In?ammasome. Cancer Res 81, 2745–2759 (2021). 10.1158/0008-5472.Can-20-3827

26 Díaz-Basabe, A. et al. Porphyromonas gingivalis fuels colorectal cancer through CHI3L1-mediated iNKT cell-driven immune evasion. Gut Microbes 16, 2388801 (2024). 10.1080/19490976.2024.2388801

27 Genomics, X. Custom Probe Design for Visium Spatial Gene Expression and Chromium Single Cell Gene Expression Flex. Technical Note 2025. < https://cdn.10xgenomics.com/image/upload/v1729114202/support-documents/CG000621_CustomProbeDesign_RevD.pdf >.

28 Johansson, M. E. V. & Hansson, G. C. in Mucins: Methods and Protocols (eds Michael A. McGuckin & David J. Thornton) 229–235 (Humana Press, 2012).

29 Nalluri-Butz, H. et al. A pilot study demonstrating the impact of surgical bowel preparation on intestinal microbiota composition following colon and rectal surgery. Scientific Reports 12, 10559 (2022). 10.1038/s41598-022-14819-1

30 Dohlman, A. B. et al. The cancer microbiome atlas: a pan-cancer comparative analysis to distinguish tissue-resident microbiota from contaminants. Cell Host & Microbe 29, 281–298.e285 (2021). 10.1016/j.chom.2020.12.001

31 Lee, S. et al. Global compositional and functional states of the human gut microbiome in health and disease. Genome Res 34, 967–978 (2024). 10.1101/gr.278637.123

