## Supplementary Table 1 for "Custom Probe-Based Spatial Transcriptomics Enables Microbiome Detection in FFPE Colorectal Cancer Tissue"

### SUPPLEMENTARY FIGURES

| Target Taxon (Probe Group) | Species of Interest | NCBI RefSeq Accession |
|---|---|---|
| **Alistipes** | *Alistipes finegoldii* | NR_103154.1 NR_115300.1 NR_102944.1 NR_043064.1 NR_113150.1 |
| | *Alistipes onderdonkii* | NR_043318.1 NR_113151.1 |
| | *Alistipes putredinis* | NR_113152.1 NR_025909.1 |
| **Bacteroides-Phocaeicola** | *Bacteroides clarus* | NR_113065.1 NR_112893.1 |
| | *Bacteroides congonensis* | NR_179552.1 |
| | *Bacteroides eggerthii* | NR_040864.1 NR_112935.1 |
| | *Bacteroides fragilis* | NR_112936.1 NR_074784.2 NR_112141.1 NR_119164.1 |
| | *Bacteroides intestinalis* | NR_041307.1 |
| | *Phocaeicola massiliensis* | NR_042745.1 NR_112938.1 |
| | *Bacteroides nordii* | NR_043017.1 NR_112939.1 |
| | *Bacteroides salyersiae* | NR_043016.1 NR_112942.1 |
| | *Bacteroides stercoris* | NR_112943.1 NR_027196.1 |
| | *Bacteroides thetaiotaomicron* | NR_076197.1 NR_074277.1 NR_112944.1 NR_112142.1 |
| | *Phocaeicola vulgatus* | NR_076384.1 NR_074515.1 NR_112946.1 NR_112143.1 |
| **Butyricimonas** | *Butyricimonas virosa* | NR_041691.1 |
| **Campylobacter** | *Campylobacter jejuni* | NR_118520.1 NR_041834.1 NR_043599.1 NR_117760.1 |
| | *Campylobacter rectus* | NR_043606.1 NR_113247.1 |
| | *Campylobacter showae* | NR_118526.1 NR_043601.1 NR_118151.1 |
| | *Campylobacter ureolyticus* | NR_117766.1 NR_118654.1 |
| **Coprobacter** | *Coprobacter fastidiosus* | NR_118316.1 |
| **Dialister** | *Dialister invisus* | NR_113355.1 NR_025680.1 |
| | *Dialister pneumosintes* | NR_026229.1 |
| **Enterobacteriaceae** | *Escherichia coli* | NR_114042.1 NR_112558.1 NR_024570.1 |
| **Fusobacterium** | *Fusobacterium animalis* | NR_113378.1 NR_117843.1 NR_117291.1 NR_026084.1 |
| | *Fusobacterium nucleatum* | NR_074412.1 NR_113198.1 NR_117287.1 NR_114702.1 |
| | *Fusobacterium polymorphum* | NR_041807.1 NR_117842.1 NR_113141.1 NR_117288.1 |
| **Helicobacter** | *Helicobacter pylori* | NR_114587.1 NR_044761.1 NR_119304.1 |
| **Intestinimonas** | *Intestinimonas butyriciproducens* | NR_118554.1 |
| **Lachnospiraceae** | *Hungatella hathewayi* | NR_036928.1 |
| | *Enterocloster asparagiforme* | NR_042200.1 |
| | *Enterocloster bolteae* | NR_113410.1 NR_025567.1 |
| | *Roseburia inulinivorans* | NR_042007.1 |
| | *Ruminococcus torques* | NR_115502.1 NR_036777.1 |
| **Leptotrichia** | *Leptotrichia trevisanii* | NR_028769.1 |
| | *Leptotrichia wadei* | NR_036844.1 NR_113229.1 |
| **Neisseria** | *Neisseria mucosa* | NR_117696.1 NR_117717.1 |
| **Parabacteroides** | *Parabacteroides distasonis* | NR_074376.1 NR_041342.1 |
| | *Parabacteroides merdae* | NR_041343.1 NR_119166.1 |
| **Parvimonas** | *Parvimonas micra* | NR_114675.1 NR_114338.1 NR_036934.1 |
| **Peptostreptococcus** | *Peptostreptococcus stomatis* | NR_043589.1 |
| **Porphyromonas** | *Porphyromonas asaccharolytica* | NR_076875.1 NR_116811.1 NR_074588.1 NR_044635.1 NR_113079.1 |
| | *Porphyromonas endodontalis* | NR_042803.1 NR_113085.1 |
| | *Porphyromonas gingivalis* | NR_114574.1 NR_114638.1 NR_074234.1 NR_040838.1 NR_113086.1 NR_119038.1 |
| **Prevotellaceae** | *Prevotella copri* | NR_040877.1 NR_113411.1 |
| | *Prevotella intermedia* | NR_114639.1 NR_116809.1 NR_113106.1 NR_113107.1 NR_026119.1 |
| | *Prevotella nigrescens* | NR_114640.1 NR_113115.1 NR_113116.1 NR_044850.1 |
| | *Prevotella stercorea* | NR_041364.1 |
| **Streptococcaceae** | *Streptococcus anginosus* | NR_121936.1 NR_118289.1 NR_118320.1 NR_041722.2 NR_117426.1 NR_112705.1 NR_118937.1 |
| | *Streptococcus equinus* | NR_114642.1 NR_113594.1 NR_042052.1 |
| | *Streptococcus gallolyticus* | NR_044904.1 |
| | *Streptococcus sanguinis* | NR_115736.1 NR_076430.1 NR_113260.1 NR_024841.1 NR_111994.1 |
| | *Streptococcus vestibularis* | NR_042777.1 NR_118942.1 |

**Supplementary Table 1. NCBI RefSeq 16S rRNA sequence accessions used for probe design.** For each target taxon, species of interest and accession number(s) used for input for the custom probe design pipeline are listed.
